# The influence factors of mean particle size and positive quality control of bacterial filtration efficiency system

**DOI:** 10.1101/2021.12.20.473569

**Authors:** Yayi Yi, Dianlong Shi

## Abstract

**Background:** The Coronavirus Disease 2019 (COVID-19) has swept the whole world with high mortality. Since aerosol transmission is the main route of transmission, wearing a mask serves as a crucial preventive measure. An important parameter to evaluate the performance of a mask is the bacteria filtration efficiency (BFE). Aerosol mean particle size (MPS) and positive quality control value are two key indexes of BFE system.

**Aim:** To study the major influence factors of the mean particle size of bacterial aerosols and positive quality control value of BFE system.

**Method and Results:** In this study, we investigated the influence of Anderson sampler, spray flow, medium thickness, and peristaltic pump flow on the MPS of bacterial aerosols and positive quality control value of BFE system, respectively. The results show that the machining accuracy of Anderson sampler has great influence on aerosol MPS and positive quality control value. With the increase of aerosol spray flow rate, the positive quality control value will increase gradually, but the effect on aerosol MPS is not a simple linear relationship. As the agar medium thickness increased, the positive quality control value and aerosol MPS increased gradually. With the increase of peristaltic pump flow, the positive quality control value increased gradually, while the aerosol MPS was basically in a downward trend. When the peristatic pump flow rate was 0.1mL/min, the spray flow rate was 7.2L/min, the agar plate thickness was 27mL, and the Anderson sampler of Beijing Mingjie was used for the experiment, the aerosol MPS and positive quality control value were both within the acceptable range and were the optimal parameters.

**Conclusions:** This study provides guidance for the manufacturers of the BFE system and improves the protective performance of masks, which is important for the human health, especially during the occurrence of viral pandemics such as “COVID-19”.

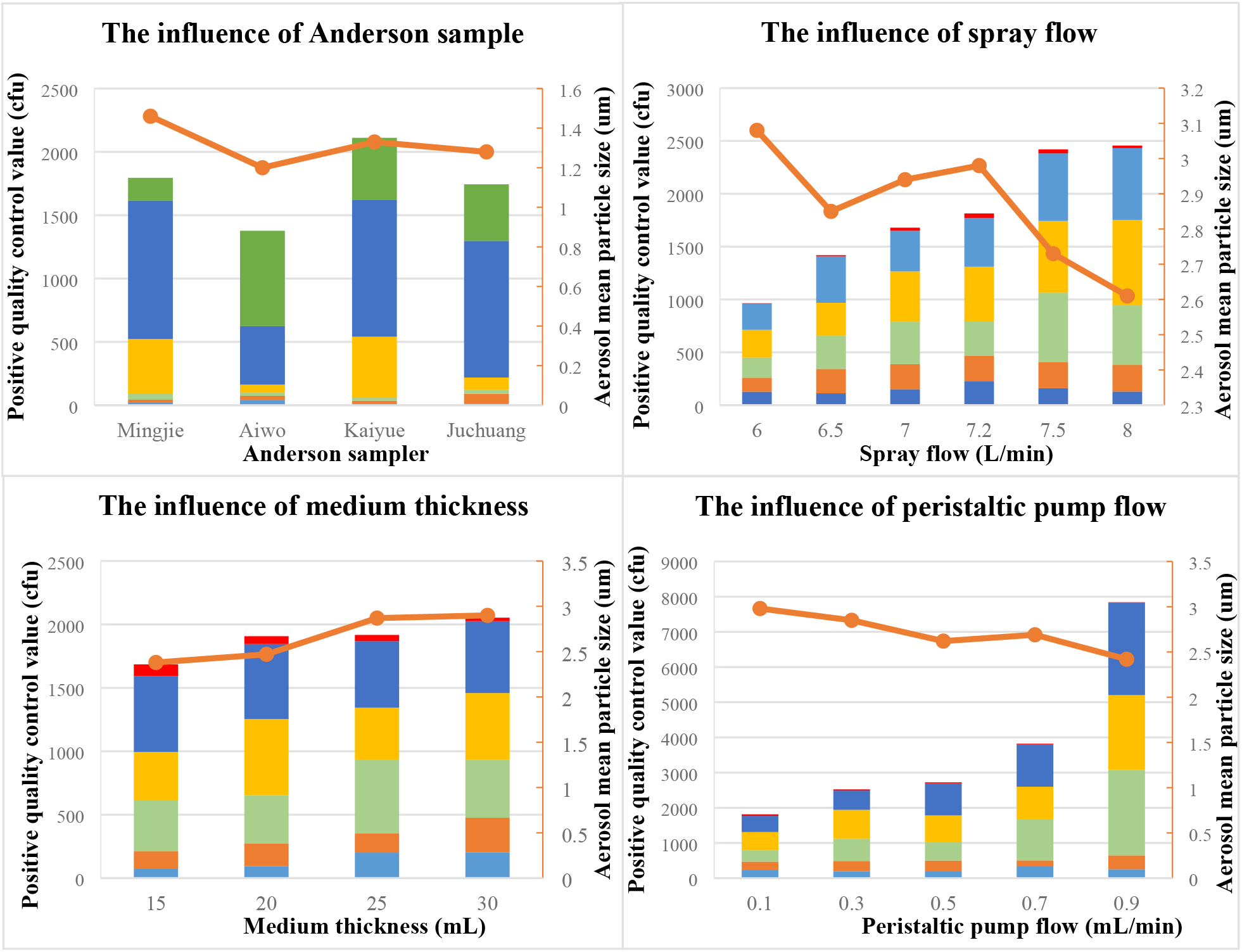

## Introduction

In December 2019, an outbreak of the Coronavirus Disease 2019 (COVID-19) occurred in the city of Wuhan, China. The COVID-19 has hit all regions in China and almost all countries worldwide at an unprecedented transmission rate, and was declared as a Public Health Emergency of International Concern (PHEIC) by the World Health Organization (WHO) ^[1]^. The occurrence of viral pandemics with extensive transmission and the associated health impact is a big concern around the globe, which raises the importance of respiratory protection against the viral transmission. Aerosol transmission is the main routes of COVID-19 transmission. Coughing, sneezing, and talking can transmits microbial aerosols through the air, potentially carrying infectious diseases ^[2]^. The associated risk can be exacerbated when pathogenic microbials including bacteria and virus are present in the air ^[3,4]^. The role of respiratory protection becomes particularly important in the occurrence of viral pandemics such as “COVID-19”, SARS, and H1N1 influenza ^[3–6]^.

Mask has been used for more than 100 years to prevent the spread of respiratory infectious diseases and surgical infections ^[7–11]^. The role of mask is to prevent harmful aerosols from the human body instead of being inhaled into the lungs, including dust, smoke, bacteria, and virus. Using masks to prevent viral transmission has been recommend by many international guidelines ^[12–14]^. MacIntyre et al. ^[15]^ reported that masks can reduce the risk of influenza infection significantly. Brienen et al. ^[16]^ showed that population-wide use of face masks could make an important contribution in delaying an influenza pandemic. Van der Sande et al. ^[17]^ have indicated that the protective factor of surgical masks was 4.1–5.3, while the protective factor of homemade masks was 2.2–2.5, which could reduce the respiratory infections of the population to a certain extent. With the outbreak of “COVID-19”, the disposable masks are in short supply, and the demand for masks bacterial filtration efficiency detector is rising rapidly ^[18]^. From 20 Jan 2020 to 30 Jun 2020, the largest daily facemask shortages in China were predicted to be 589.5, 49.3, and 37.5 million in the mask-wearing policy in all regions of mainland, the mask-wearing policy only in Hubei province, and the non-implementation of the mask-wearing policy in other region, respectively ^[19, 20]^.

An important parameter to evaluate the performance of a mask is the bacteria filtration efficiency (BFE) ^[21]^. BFE refers to the percentage of the respirator material filtered out of the bacteria-containing suspended particles under the specified flow rate, which reflects the filtration quality of the respirator. The protective performance of a mask mainly depends on the filtration efficiency of the mask. How to ensure the mean particle size (MPS) of bacterial aerosols and the positive quality control value are the most critical indicators of the mask BFE detector, and is also the current research hotspots ^[22]^. In this study, the major influence factors of the MPS and positive quality control value of the BFE system were investigated, including Anderson sampler, spray flow, medium thickness, and peristaltic pump flow, which provides guidance for the manufacturers of the BFE detector and improve the protective performance of masks under the pandemics of COVID-19.

## Materials and methods

### Materials

Tryptic soy agar (TSA), tryptic soy broth (TSB), peptone water, staphylococcus aureus ATCC6538, petri dish (φ 90 mm), and vaccination ring.

### Equipment

High pressure steam sterilization pot (121°C~123°C), incubator (constant temperature 37°C±2°C), vortex type vortex mixer (up to 16 mm × 150 mm tube), orbital vibrator (100 r/min~250 r/min), refrigerator (2°C~8°C), bacteria filtration efficiency detector (ZR-1000, Qindao Junray Intelligent Instrument Co., Ltd), and colony counter (ZR-1100, Qindao Junray Intelligent Instrument Co., Ltd).

### Experimental methods

#### Determination method of bacterial suspension concentration

A strain of staphylococcus aureus was vaccinated into 100mL TSB, then exposed to an incubator under 100r/min at 37°C. After 24h, the bacterial suspension was obtained. 1mL bacterial suspension was added to the test tube with 9mL peptone solution (1.5%), then diluted step by step to 10^−7^, and 0.1mL diluted bacterial suspension from 10^−5^ to 10^−7^ test tubes was added to TSA plate for culture. Two parallel plates were made for each gradient, then put them into the incubator at 37°C for 24h.

Statistic the bacterial count of each TSA plate, then take the colony number of (0-100) CFU flat gradient to count, and the concentration of bacteria suspension concentrate can be calculated. Then diluted 1 mL bacteria suspension to about 5 × 10^5^ CFU/mL, and 20 mL of which were prepared.

#### Positive quality control value calculation

Perform a positive control run without a test specimen by BFE detector. Initiate the bacterial suspension by turning on the vacuum pump and adjust the flow rate through the Anderson sampler to 28.3L/min. Deliver the bacterial suspension by the peristaltic pump for 1 min. Sampling for 2 min, and the bacteria aerosols were collected on agar medium. Agar plate were cultivated in the incubator at (37±2)°C for (24±4) h, then count the number of staphylococcus aureus colonies on each plate, and use the “positive hole” conversion table in accordance with the instructions of the Anderson sampler manufacturer for stages 3 to 6. The sum of staphylococcus aureus colonies on each stage is a positive quality control value of detector, and take the mean of the two totals for two positive control runs. ASTM F2101-2014 Evaluating the bacterial filtration efficiency (BFE) of medical face mask materials, using a biological aerosol of staphylococcus aureus ^[23]^ and BS EN 14683:2019 Medical face masks - Requirements and test methods ^[24]^ specifies, positive quality control value should be within the scope of the (1700-3000) CFU.

#### Bacterial aerosol MPS calculation

After sampling, the agar medium were cultivated in the incubator at (37±2)°C for (24±4) h. From the positive control plates, the MPS of bacterial aerosol was calculated using the formula (1). BS EN 14683:2019 Medical face masks - Requirements and test methods ^[24]^ and YY0469-2011 Medical surgical masks technical requirements ^[25]^ rules, the MPS should be within the scope of (3.0±0.3) um.

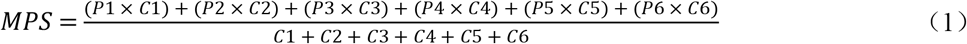

## Results and discussion

### The influence of Anderson sampler

Four Anderson sampler from different manufacturers were selected to compare the influence of Anderson sampler in the aerosol MPS and positive quality control value, including Beijing Mingjie, Qingdao Kaiyue, Qingdao Juchuang and Weifang Aiwo. The experiment condition was set as: the peristaltic pump flow is 0.1 mL/min, the spray flow is 10 L/min, and agar medium thickness is 15 mL. Positive quality control values and aerosol MPS with different Anderson sampler manufacturer were shown in Table 2.

**Table 1.** Anderson sampler particle size and colony count at all levels.

|  |  |  |  |  |  |  |
| --- | --- | --- | --- | --- | --- | --- |
| Stage number | 1 | 2 | 3 | 4 | 5 | 6 |
| Size of particle | P1 | P2 | P3 | P4 | P5 | P6 |
| Viable “particle” plate count | C1 | C2 | C3 | C4 | C5 | C6 |

Where P1=7.00um ;P2=4.70um ;P3=3.30um ;P4=2.10um ;P5=1.10um ;P6=0.65um.
C1+C2+C3+C4+C5+C6 refer to positive quality control value.

**Table 2.** Positive quality control values and aerosol MPS with different Anderson sampler manufacturer.

| Sampler | Beijing Mingjie |  | Weifang Aiwo |  |
| --- | --- | --- | --- | --- |
| Peristaltic pump flow | 0.1 mL/min |  | 0.1 mL/min |  |
| Spray flow | 10 L/min |  | 10 L/min |  |
| Spray pressure | 21.2 kPa |  | 21.2 kPa |  |
| Tablet thickness | 15 mL |  | 15 mL |  |
| The layer number | The positive pole count (r) | Viable “particle” plate count (C) | The positive pole count (r) | Viable “particle” plate count (C) |
| 1 | 19 | 19 | 40 | 40 |
| 2 | 25 | 25 | 34 | 34 |
| 3 | 43 | 45 | 23 | 24 |
| 4 | 265 | 434 | 60 | 65 |
| 5 | 374 | 1093 | 274 | 462 |
| 6 | 144 | 179 | 339 | 752 |
| Positive quality control value | / | 1795 | / | 1377 |
| MPS |  | 1.46 |  | 1.20 |

| Sampler | Qingdao Kaiyue |  | Qingdao Juchuang |  |
| --- | --- | --- | --- | --- |
| Peristaltic pump flow | 0.1 mL/min |  | 0.1 mL/min |  |
| Spray flow | 10 L/min |  | 10 L/min |  |
| Spray pressure | 20.8 kPa |  | 20.8 kPa |  |
| Tablet thickness | 15 mL |  | 15 mL |  |
| The layer number | The positive pole count (r) | Viable “particle” plate count (C) | The positive pole count (r) | Viable “particle” plate count (C) |
| 1 | 13 | 13 | 10 | 10 |
| 2 | 23 | 23 | 81 | 81 |
| 3 | 22 | 23 | 28 | 29 |
| 4 | 280 | 482 | 88 | 99 |
| 5 | 370 | 1078 | 373 | 1078 |
| 6 | 283 | 492 | 269 | 447 |
| Positive quality control value | / | 2069 | / | 1744 |
| MPS |  | 1.33 |  | 1.28 |

By comparing and testing the Anderson sampler from different manufacturers, it is found that the aerosol MPS and positive quality control value calculated from the experimental results of the Anderson sampler from different manufacturers is different under the same experimental conditions. The positive quality control value of other three experiments were all within the scope of (1700-3000) CFU except using the Weifang Aiwo’s Anderson sampler. The order of aerosol MPS is Beijing Mingjie > Qingdao Kaiyue > Qingdao Juchuang > Weifang Aiwo, so the Beijing Mingjie’ Anderson sampler was recommended. The design principle of Anderson sampler is as follows: when microbial aerosols pass through a nozzle or jet stream, they shoot at the front impingement plate (or the surface of the medium) according to the principle of inertia, deflecting the airflow 90°~180°. Particles of sufficient momentum, acting by inertia, move in a straight line in the original direction without following the deflection of the fluid, and are collected by impingement on the collecting plate (or on the surface of the medium). Smaller particles can follow the fluid along the streamline under the air flow without crashing down, and these bacteria-bearing particles will slip off or escape because of their small inertia. The Anderson sampler can divide the bacterial aerosol into two parts. Particles larger than a certain aerodynamic diameter can crash down from the airflow, and smaller particles can escape with the aerosol fluid passing through the sampler. As each manufacturer has different requirements for pore accuracy, the size of bacterial aerosol captured by Anderson sampler of each manufacturer is also different, so the calculated aerosol MPS is also different. The pore size and capture particles range of Anderson sampler is shown in Table 3.

**Table 3.** The pore size and scope of capture particles of Anderson sampler.

| Anderson sampler stages | Pore size (mm) | The scope of capture particles (um) |
| --- | --- | --- |
| Stage 1 | 1.18 | >7.0 |
| Stage 2 | 0.91 | 4.7~7.0 |
| Stage 3 | 0.71 | 3.3~4.7 |
| Stage 4 | 0.53 | 2.1~3.3 |
| Stage 5 | 0.34 | 1.1~2.1 |
| Stage 6 | 0.25 | 0.65~1.1 |

### The influence of spray flow rate

Different spray flow rate was set to compare the effects of spray flow in the aerosol MPS and positive quality control value, including 6.0, 6.5, 7.0, 7.2, 7.5, and 8.0 L/min. The experiment condition was set as: the peristaltic pump flow is 0.1 mL/min, agar medium thickness is 25 mL, and Beijing Mingjie’ Anderson sampler was selected.

The positive quality control values and aerosol MPS at different spray flow rates were shown in Table 4. The positive quality control value increases gradually with the increase of the flow rate of the sprayer, which may due to the increasing spray flow blowing more bacterial aerosols to the air chamber. However, there is no linear relationship between aerosol MPS and aerosol flow rate (Figure 3). When the spray flow rate is 7.2 L/min and 7.5 L/min, the positive quality control values conform to the range of (1700-3000) CFU, and the MPS of aerosol fall within the scope of (3.0±0.3) um. Compared to the MPS of aerosol under spray flow rate of 7.5 L/min, when the spray flow rate is 7.2 L/min, the MPS is maximum and the flow for this sprayer is the most appropriate option on the premise of meeting the standard value of positive quality control. The mean aerosol particles depend on comprehensive influence, including the size of the sprayer, spray flow rate, flow of peristaltic pump, physical properties of bacteria aerosol, pore diameter, the relative position and angle of fluid pore in mixing chamber of sprayer, so the appropriate spray flow will be different for every sprayer due to very slight error on each sprayer processing.

**Table 4.** Positive quality control values and aerosol MPS at different spray flow rates.

| Number | Spray flow<br>(L /min) | Positive quality control<br>value (CFU/mL) | Aerosol MPS (um) |
| --- | --- | --- | --- |
| 1 | 6.0 | 964 | 3.08 |
| 2 | 6.5 | 1419 | 2.85 |
| 3 | 7.0 | 1679 | 2.94 |
| 4 | 7.2 | 1814 | 2.98 |
| 5 | 7.5 | 2420 | 2.73 |
| 6 | 8.0 | 2457 | 2.61 |

**Figure 1.**
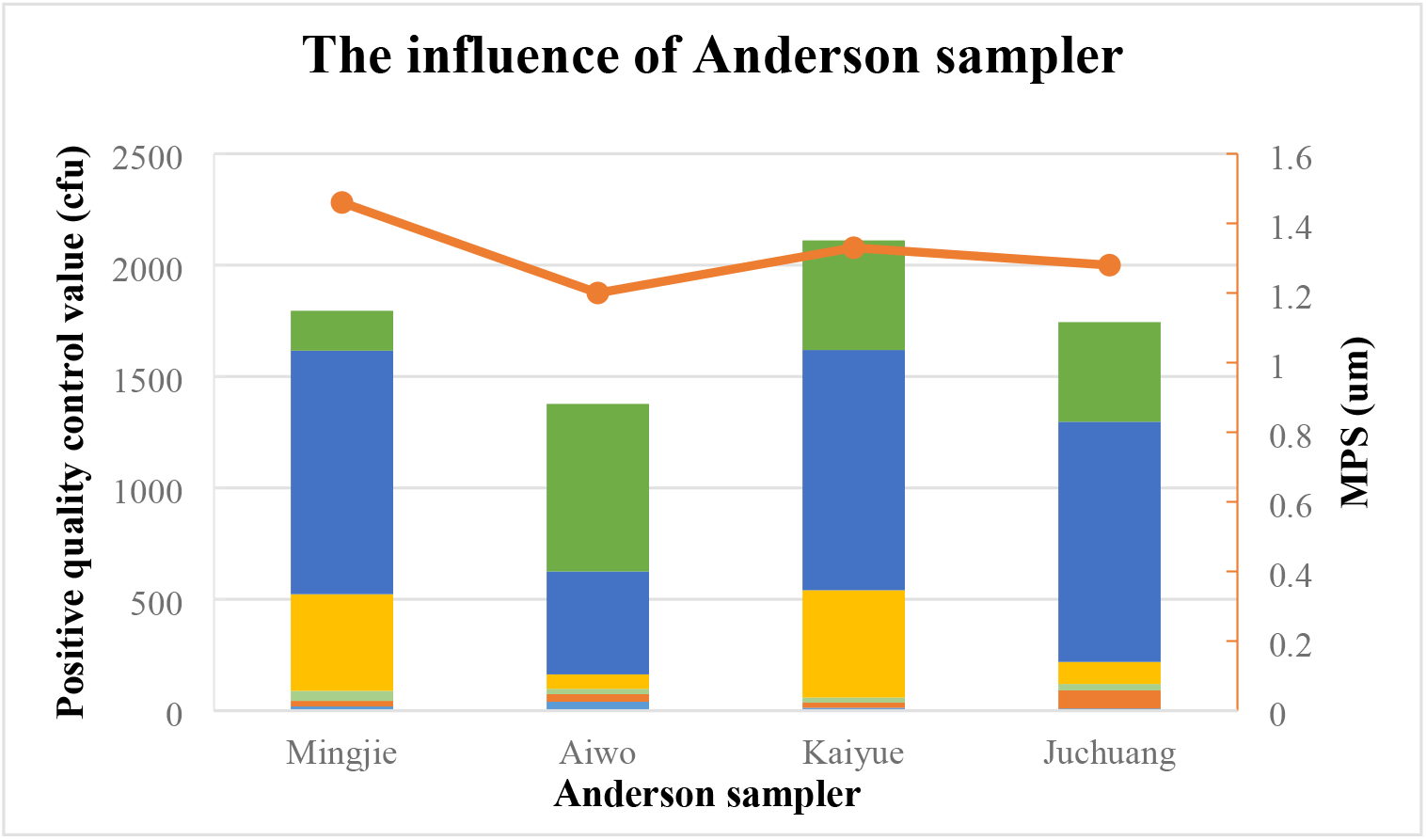
The relationship of Anderson sampler with positive quality control value and aerosol MPS (Histogram represents positive quality control value, broken line represents aerosol MPS).

**Figure 2.**
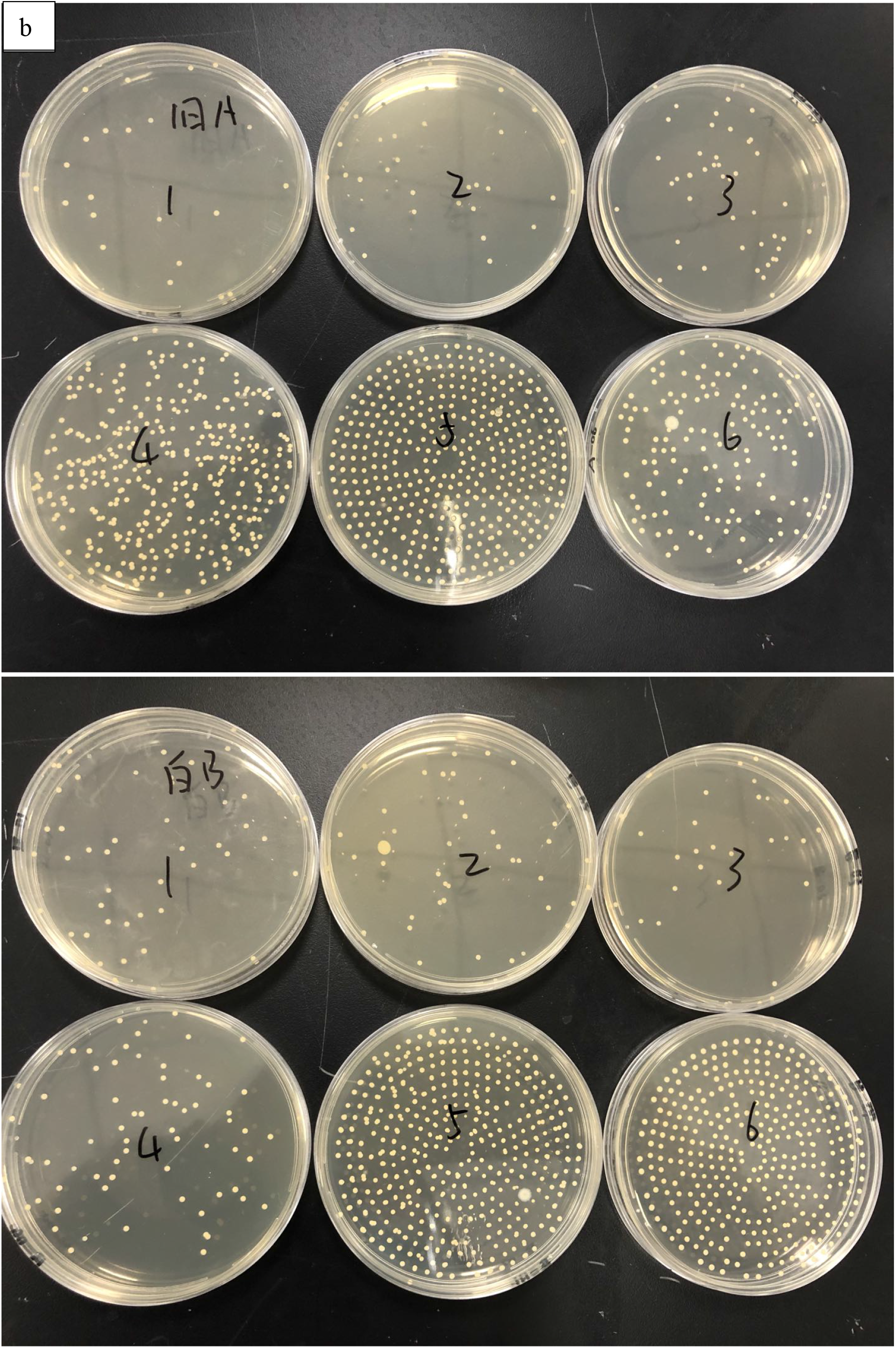
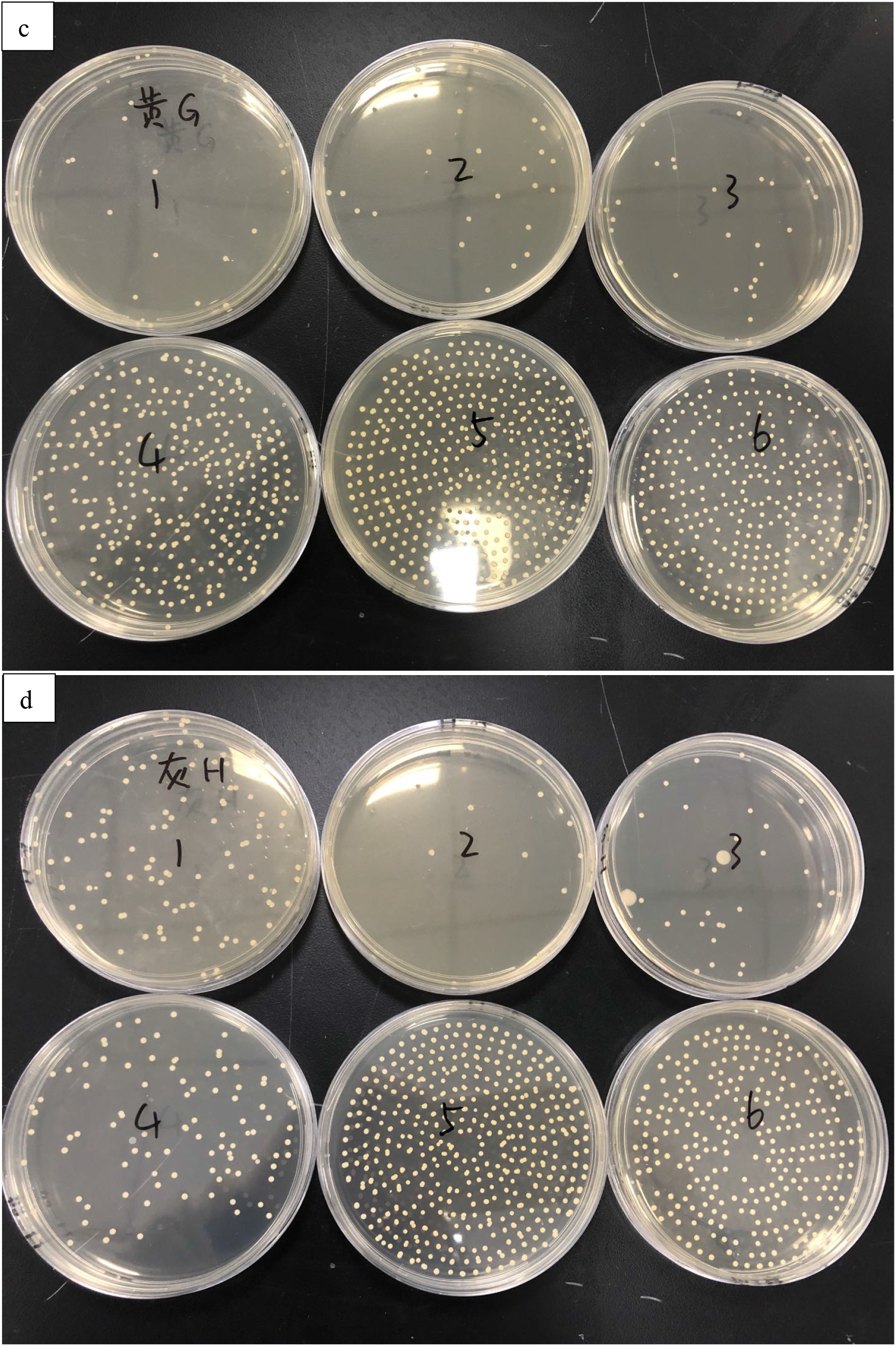
The experimental results of different Anderson sampler (a for Beijing Mingjie, b for Weifang Aiwo, c for Qingdao Kaiyue, d for Qingdao Juchuang).

**Figure 3.**
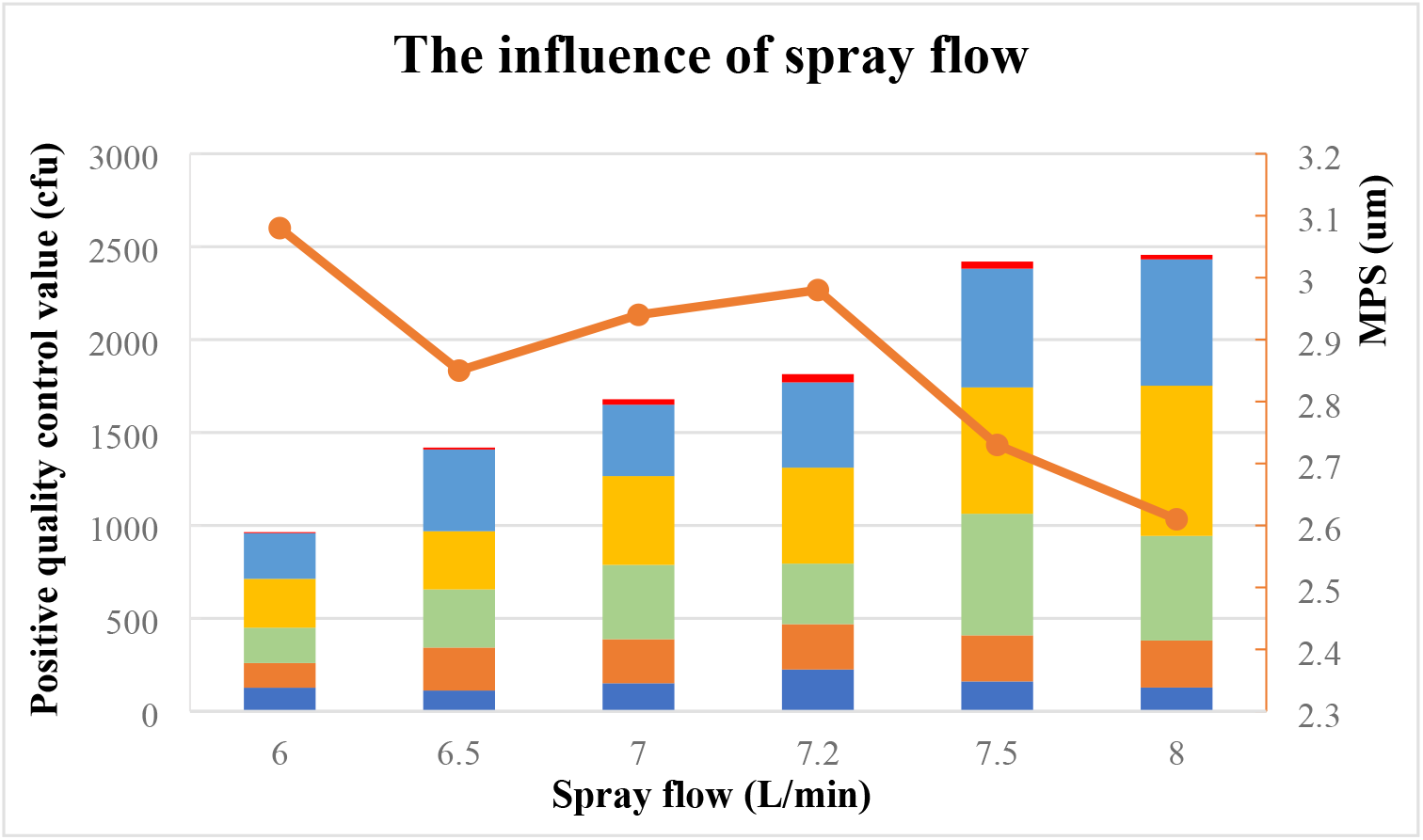
The relationship of spray flow with positive quality control value and aerosol MPS (Histogram represents positive quality control value, broken line represents aerosol MPS).

### The influence of agar medium thickness

Different agar medium thickness of 15, 20, 25, and 30 mL were prepared to compare the influence of the thickness of agar medium in the aerosol MPS and positive quality control value. The experiment condition was set as: the peristaltic pump flow is 0.1 mL/min, the spray flow rate is 7.2 L/min, and Beijing Mingjie’ Anderson sampler was selected.

The positive quality control values and aerosol MPS under different medium thickness were shown in Table 5. As the agar medium thickness increased, both the positive quality control value and aerosol MPS increased gradually (Figure 4). When the agar medium thickness was 25mL or 30mL, the MPS could meet the requirement of (3±0.3) um. This is because the impact distance has an influence on the sampling efficiency. The aerosol particles cannot come down if the distance is too far, while it is difficult to operate with short distance. The impact distance of the Anderson sampler is about 2.5mm. When the sampler structure is fixed, the thickness of the agar medium in the plate determines the impact distance. Adding 24, 27 and 30mL agar medium to the petri dish, respectively, and their impact distances were 3, 2.5 and 2.0mm. But the percentage distribution of the three impact distance particles in the six nodes was roughly similar. Therefore, Ranz and Wong ^[26]^ recommend that adding 27mL agar medium into Anderson sampler is optimal.

**Table 5.** Positive quality control values and aerosol MPS under different medium thickness.

| Number | Medium thickness<br>(mL) | Positive quality control<br>value (CFU/mL) | Aerosol MPS (um) |
| --- | --- | --- | --- |
| 1 | 15 | 1685 | 2.38 |
| 2 | 20 | 1907 | 2.47 |
| 3 | 25 | 1917 | 2.87 |
| 4 | 30 | 2053 | 2.90 |

**Figure 4.**
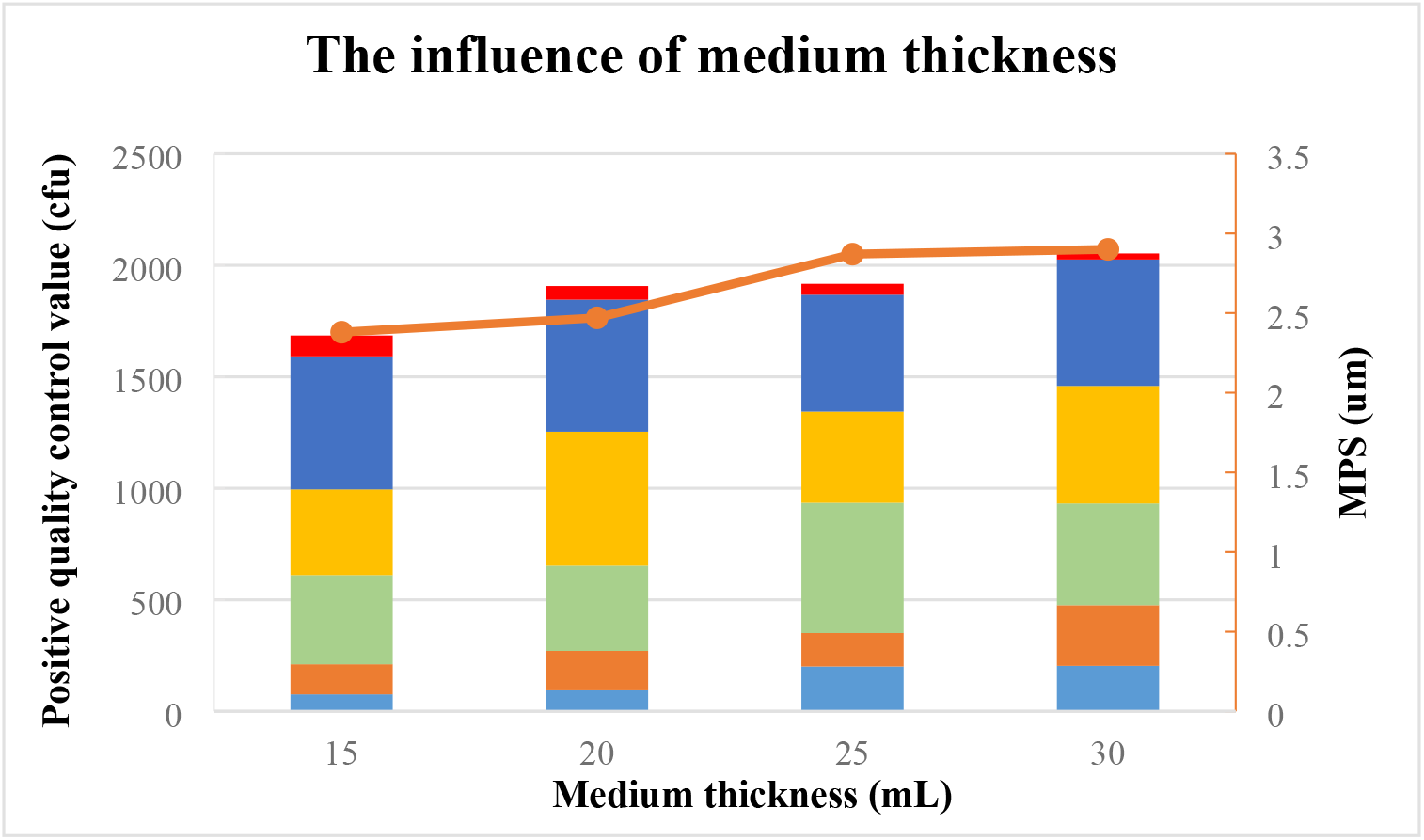
The relationship of medium thickness with positive quality control value and aerosol MPS (Histogram represents positive quality control value, broken line represents aerosol MPS).

### The influence of peristaltic pump flow

Different peristaltic pump flow rate was selected to compare the influence of peristaltic pump flow in the aerosol MPS and positive quality control value, including 0.1, 0.3, 0.5, 0.7, 0.9 mL/min. The experiment condition was set as: the spray flow rate is 7.2 L/min, agar medium thickness is 25 mL, and Beijing Mingjie’ Anderson sampler was selected.

The positive quality control values and aerosol MPS under different peristaltic pump flow were shown in Table 6. With the increase of peristaltic pump flow, the positive quality control value gradually increased, while the aerosol MPS was basically in a downward trend (Figure 5). When the peristaltic pump flow was 0.1mL/min, the aerosol MPS reached its maximum value. This may be attributed to that the microbial quantity of bacterial aerosols entering the sprayer increased with the peristaltic pump flow under the same spray flow. However, small particle aerosols produced far more than large particles of bacterial aerosols, so the MPS of aerosols in a downtrend basically although the positive quality value increases gradually. Therefore, we suggest that the peristaltic pump flow of 0.1mL/min is the most appropriate option.

**Table 6.** Positive quality control values and aerosol MPS under different peristaltic pump flow.

| Number | Peristaltic pump flow<br>(mL /min) | Positive quality control<br>value (CFU/mL) | Aerosol MPS (um) |
| --- | --- | --- | --- |
| 1 | 0.1 | 1814 | 2.98 |
| 2 | 0.3 | 2523 | 2.85 |
| 3 | 0.5 | 2723 | 2.62 |
| 4 | 0.7 | 3821 | 2.69 |
| 5 | 0.9 | 7842 | 2.42 |

**Figure 5.**
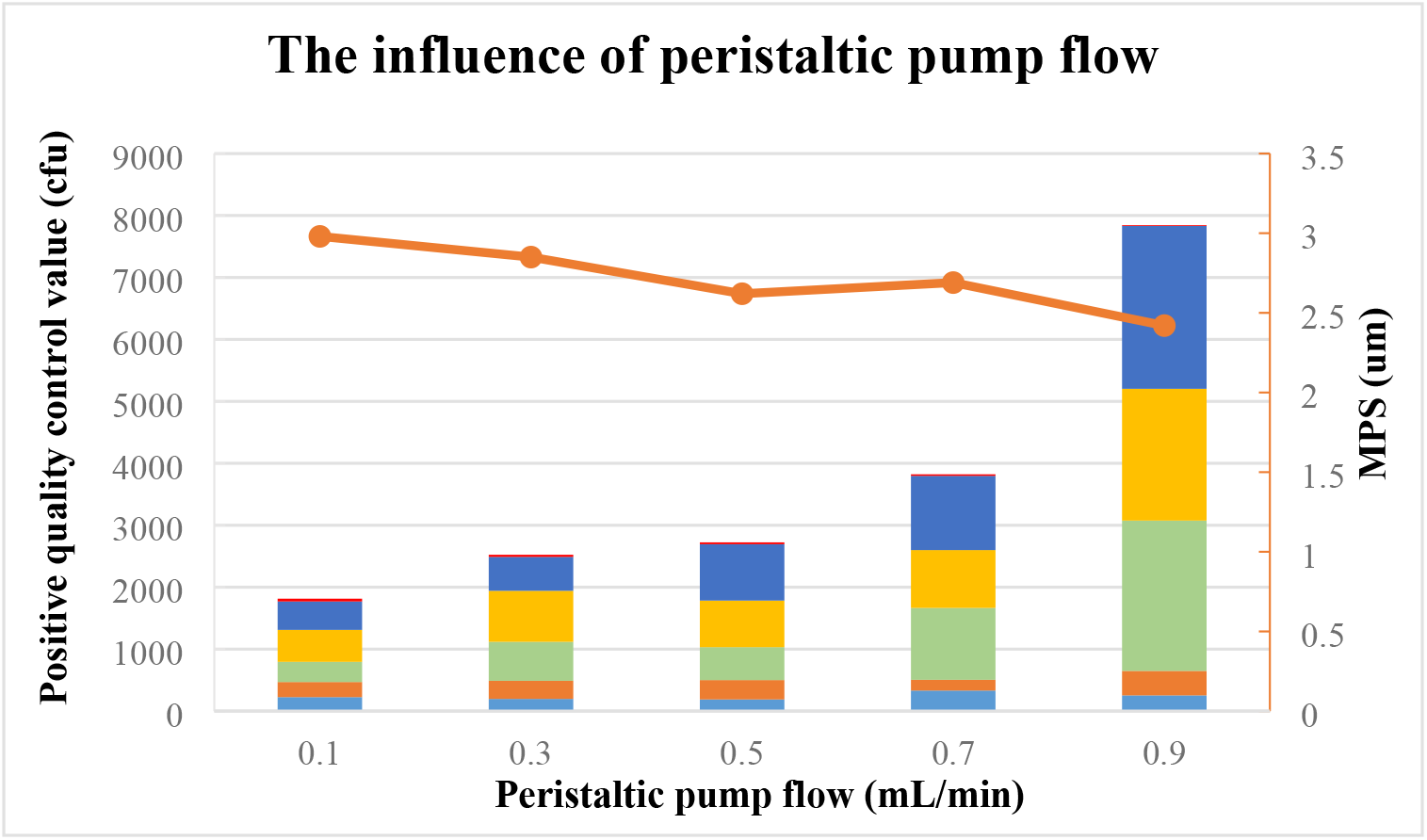
The relationship of peristaltic pump flow with positive quality control value and aerosol MPS (Histogram represents positive quality control value, broken line represents aerosol MPS).

## Conclusion

The mask BFE is one of the key indicators for detecting masks. The aerosol MPS and positive quality control value are the key parameters of BFE system. This study discusses the major influence factors of the MPS of bacterial aerosols and positive quality control value of BFE system, including Anderson sampler, spray flow, agar medium thickness, and peristaltic pump flow. The experimental results show that the machining precision of Anderson sampler have great influence on aerosol MPS and positive quality control value. With the increase of spray flow rate, the positive quality control value will increase gradually, but the effect on aerosol MPS is not a simple linear relationship. As agar medium thickness increased, the positive quality control value and aerosol MPS increased gradually. With the increase of peristaltic pump flow, the positive quality control value increased gradually, while the aerosol MPS was basically in a downward trend. When the peristatic pump flow rate was 0.1mL/min, the spray flow rate was 7.2L/min, the agar medium thickness was 27mL, and the Anderson sampler of Beijing Mingjie was used for the experiment, the positive quality control value and aerosol MPS were both within the acceptable range and were the optimal parameters. This study provides guidance for manufacturers of BFE detector and theoretical basis for testing institutions to test the BFE of masks, which is important for the human health, especially during the occurrence of viral pandemics such as “COVID-19”.

## Notes

### Competing Interest Statement

The authors have declared that no competing interests exist.

